# Removing EOG Artifacts from EEG Recordings using Deep Learning

**DOI:** 10.1101/2024.05.24.595839

**Authors:** C. O’Reilly, S. Huberty

## Abstract

The electroencephalogram (EEG) directly measures the electrical activity generated by the brain. Unfortunately, it is often contaminated by various artifacts, notably those caused by eye movements and blinks (EOG artifacts). Such artifacts are usually removed using an independent component analysis (ICA) or other blind source separation techniques. However, it is difficult to assess whether subtracting EOG components estimated through ICA removes some neurogenic activity. It is crucial to address this question to avoid biasing EEG analyses. Toward that objective, we developed a deep learning model for EOG artifact removal that exploits information about eye movements available through eye-tracking (ET). Using a multimodal EEG and ET open-access dataset, we trained within-subject a long short-term memory (LSTM) model to predict the component of EEG signals predictable from ET data. We further used this ET-informed evaluation of EOG artifacts to investigate the sensitivity and specificity of ICA. Our analysis indicates that although ICA is very sensitive to EOG, it has a comparatively low specificity. These results motivate further research on EEG artifact removal to develop approaches with higher EOG rejection specificity.

## 1. Introduction

Electroencephalography (EEG) is a non-invasive neuroimaging technique used to record the electrical activity generated by the brain. EOG (electrooculogram) artifacts in EEG recordings refer to the electrical signals generated by eye movements and eye blinks due to the potential difference between the cornea and the retina, which acts as an electric dipole. When the eyes move, their dipoles also move, generating a change in the field of electric currents propagating instantaneously (quasistatic approximation [1]) through the volume of the head and reaching the EEG electrodes [2]. Thus, EOG artifacts are omnipresent in EEG signals. They usually account for much of the EEG variance, particularly in the channels near the eye. These artifacts are particularly problematic in EEG tasks requiring eye movements, as they can obscure the neural activity related to the experimental task. Therefore, developing techniques for removing EOG artifacts with high specificity is critical for EEG research, particularly for analyses of frontal connectivity involving non-lagged homotopic synchronization, which cannot be reliably distinguished from instantaneous electrical volume conduction [3].

Various techniques have been proposed to separate and remove the artifactual components from the neural components of the EEG signals. Among that class of algorithms, Independent Component Analysis (ICA) has become arguably the most widely adopted approach [4] and is used in most EEG preprocessing pipelines [5], [6]. However, although this approach has the benefit of retaining the complete EEG time course, it may fail to remove all the artifacts (insufficient sensitivity) or may distort the neural signals (insufficient specificity). Since ICA is an unsupervised algorithm, confirming that the components labeled as artifacts do not include neural signals is challenging, and some neural signals can inadvertently and unknowingly be lost when these components are subtracted from the EEG data.

While ICA can be used to identify and remove various artifacts (EOG, ECG, power line noise), alternative artifact reduction techniques for removing EOG artifacts specifically exist. For example, EOG channels from electrodes placed near the eyes can be used in a linear regression to estimate the corresponding EOG artifact in EEG channels, which can then be subtracted from the signal [7], [8]. However, EOG electrodes differ from EEG electrodes only in their placement near the eyes. Thus, they also pick up neural signals or other types of artifacts (e.g., muscle contraction, sweating), which diminishes their utility as a reference signal for EOG activity.

In this study, we aim to use deep learning to remove EOG artifacts in an EEG/eye-tracking (ET) dataset and compare the performance of this approach to ICA, the most prominent approach for EOG removal.

## 2. Methods

### 2.1. Datasets

To develop and test the proposed model, we leveraged the EEGEyeNet dataset [9]. This dataset contains recordings from 356 healthy adults, including simultaneously collected high-density 129-channel EEG data synchronized with video-infrared ET. The ET data include two channels for the position in X and Y and one for the pupil size. Both raw and preprocessed data are available in the EEGEyeNet dataset. The experiment contains three tasks: a pro- and antisaccade task, a visual symbol search task, and the *Large Grid* task. For our study, we used the latter [10]. For this task, 30 participants were asked to look at dots appearing at 25 different positions, distributed across the whole surface of a screen (see Fig. 1). Each dot is presented for 1.5 to 1.8 seconds, in a pseudo-randomized order (see [9] for details on this pseudo-randomization). The central dot was displayed three times, while the other dots were presented once per block. Each of the six runs contained five blocks, totaling 810 stimuli per participant. Each run is saved as a separate recording, providing 177 recordings (i.e., three were missing).

**Fig. 1.**
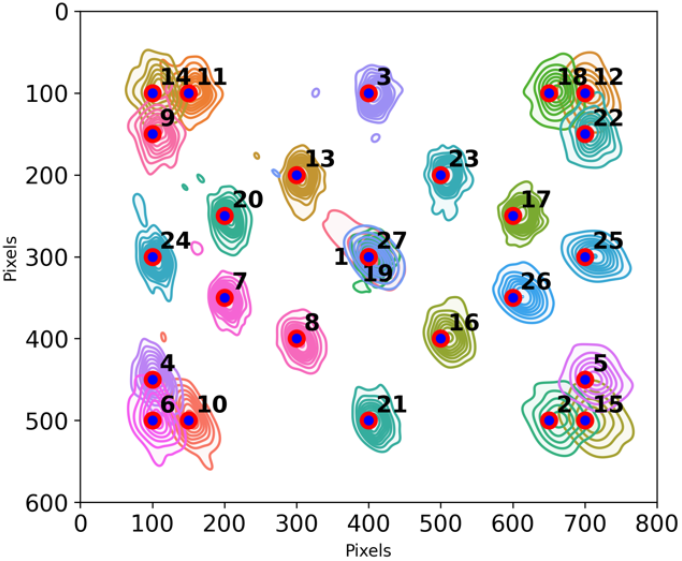
Positions of presented dots during the Large Grid task (blue/red circles) overlayed with the gaze distribution.

### 2.2. Outlier rejection

Before running analyses, we excluded recordings with noisy or unreliable ET data, which could have been due to many reasons, the most likely being poor eyetracker calibration. We identified outliers by first computing the mean squared difference between the event-related response (ERR) of every recording and the ERR averaged across recordings

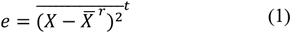

where *X* is a matrix of *x* and *y* gaze coordinates, and the bars with *r* and *t* represent the averaging across the recording and time dimensions, respectively. We computed these errors for each ET channel, dot stimuli, and recording. We then computed the 25th, 50th, and 75th quantiles of the distribution of these errors. To detect outliers in individual recordings, we used the classic outlier rejecformula, *e* > *Q*_50_ + *k*(*Q*_75_ – *Q*_50_) with *k* = 6, and rewrote

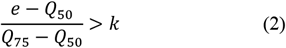

to average the left-hand term across channels and dots (i.e., event types) before comparing these averages against the threshold. Using this outlier criterion, we excluded eight recordings from further analyses. Most of them were for different runs from the same participant.

We confirmed participant compliance with the Large Grid Task instructions and the quality of the ET data for the remaining recordings by displaying the two-dimensional kernel density estimation of the distribution of the X/Y pixel coordinates for every dot in the large grid (Fig 1). For this computation, we determined the gaze position as the average position in the *t* ∈ [0.3, 1.0] s time window, with *t* = 0 being the stimulus presentation onset.

### 2.3. Preprocessing

*Minimally* and *maximally* preprocessed versions of the EEGEyeNet dataset are available [11]. These two alternative preprocessing are defined by the *Automagic* toolbox [12]. We used the minimally preprocessed version, which includes bad channel detection and interpolation and EEG filtering to the 0.5-40 Hz band. This minimal preprocessing does not include ICA artifact rejection since this step would remove the EOG artifacts necessary for our study. The authors of EEGEyeNet synchronized the EEG and ET signals and confirmed the absence of synchronization errors exceeding 2 ms.

To make our analysis more computationally efficient, we filtered to the 1-30 Hz band before downsampling the signals to 100 Hz using MNE-Python [13]. Although the dataset is recorded with a sampling rate of 500 Hz, EOG signals are limited to relatively low frequencies due to natural biomechanical constraints imposed on the kinematics of eye movements. Thus, such a high sampling rate significantly increases the network size (i.e., multiply by a factor of five the long short-term memory (LSTM) input shape and the associated weights to learn) without adding relevant information.

For machine learning, recordings were epoched into contiguous 1 s segments (this should not be confused with the concept of training epochs in deep learning), resulting in a 3D matrix of size n_segments_ × n_channels_ × n_times_. We also set an average reference. For our comparison with ICA, we used the Extended Infomax approach [14] as implemented in MNE-Python. EOG-associated components were detected automatically as those labeled as *eye blink* by MNE-ICALabel [15], which ports to Python the functionalities of ICLabel [16]. ICLabel has six additional classes of independent components: *brain, muscle artifact, heartbeat, line noise, channel noise*, and *other*.

### 2.4. Deep learning model

The overarching idea of our approach is to train a recurrent neural network (RNN) to predict EEG signals only from ET signals. Of course, only a small portion of the EEG signals will be predictable from ET signals, and this predictable portion will be due to EOG artifacts and potentially some neural and non-neural correlates of eye movements (e.g., electromyographic signals due to the activation of the muscle required for eye movements). More formally, for the EEG signal matrix *Y* (EEG channels × time) and the ET matrix *X* (ET channels × time), we model this relationship as *Y* = *f*(*X*) + *R* where *R* is a residual matrix containing the neural signals and potentially non-eye-movement-related artifacts, and *f* is a nonlinear function we want to learn by adjusting the RNN weights to minimize the mean square amplitude of *R*. For this task, we used a 2-layer LSTM with three features corresponding to ET channels and 64 hidden states whose outputs get pruned with a 0.5 dropout layer. A final fully connected layer maps the LSTM internal states to 129 outputs corresponding to the EEG channels (Fig. 2). We implemented this model in PyTorch and fitted it using the ADAM optimizer with a 0.01 learning rate and a mean-square-error (MSE) loss function. We found that 1000 training epochs (not to be confused with the EEG 1 s epochs) were enough to reach a stable training loss. We did not attempt to test for generalizability across participants or recordings. Rather, we wanted to test if the mapping between ET signals and their impact on EEG was learnable within participants. Thus, we implemented no hold-out or cross-validation. That is, the mapping was learned independently for all 177 recordings, and we used each of these mappings to clean the EEG of the corresponding recording only.

**Fig. 2.**
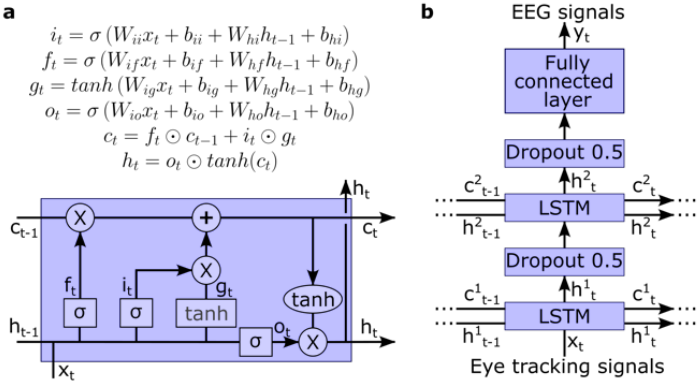
The deep neural network we designed for EOG/EEG decounfounding. **a)** Block diagram and corresponding equations for the LSTM model used in the deep neural network. In the equations, ⊙ stands for the Hadamard product, σ is the sigmoid function, lowercase letters are vectors, and uppercase letters are matrices. The matrices *W* and vectors *b* are learned during the training. **b)** Deep neural network architecture using two LSTM layers and one fully connected layer.

### 2.5. Analysis

EOG signals are known to affect mostly frontal EEG channels. To validate the capability of the neural network to detect and remove EOG noise, we computed the percentage of signal removed per channel as

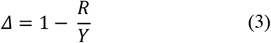

We also defined the reaction time (RT) to a stimulus as the moment the gaze position changed by 5% of maximal response amplitude. Using this RT, we used an approach similar to the one adopted in [17] to characterize the sensitivity and specificity of removing eye artifacts based on ET versus automated ICA. In this approach, we defined a pre-reaction time (pre-RT) segment where we expect no noise, here defined as the window from the start of the baseline period (−0.2 s) to the reaction time (RT), and a post-RT window where EOG artifacts are expected (RT to 1 s). Defining the signal (*S*) as the root mean square (RMS) amplitude of the original recording (*Y*) and the noise (*N*) as the RMS amplitude of what has been removed by the artifact removal approach (i.e., *N* = *Y* − *R*), we can define the classic measure of signal-to-noise ratio (SNR) in dB as

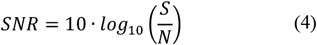

A large SNR before the reaction time is indicative of a high specificity (i.e., clean signals do not get distorted by artifact removal), and a low SNR after the reaction time is indicative of a good sensitivity (i.e., more noise has been detected and removed by the cleaning approach).

## 3. Results

A demonstration of cleaned EEG signals using the proposed model and ICA compared to the original EEG signals is shown in Fig. 3.

**Fig. 3.**
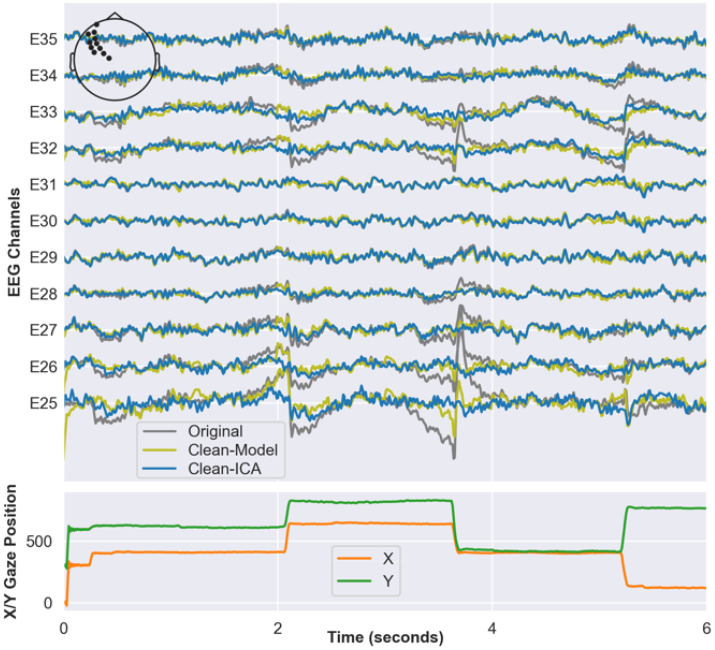
Top panel: EEG signals for a typical recording before (gray) and after EOG artifact reduction using the proposed model (green) and ICA (blue). Bottom panel: X and Y gaze positions (reported in pixel coordinates) during the same period as the top panel.

Further, evoked responses to gazes at dots confirm that the model accurately removes the EOG contamination in the EEG recordings (Fig. 4).

**Fig. 4.**
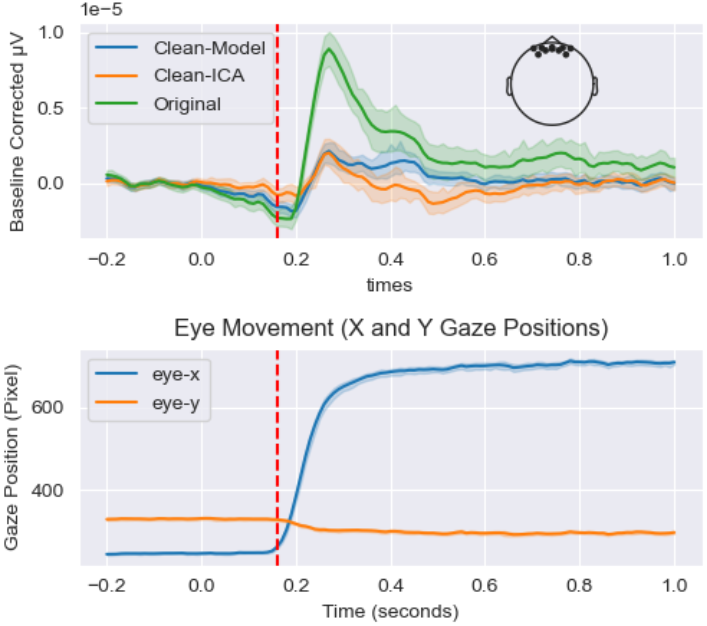
The evoked response to dot 25 before and after EOG removal (top panel) and the average eye movement (X/Y pixels) during those trials (bottom panel). The vertical red dashed line represents the average RT across participants, corresponding to the moment they began shifting their gaze toward the dot displayed at the beginning of the trial (i.e., at *t* = 0 s).

We also looked at the distribution of the predicted noise over the scalp (Fig. 5), averaged across participants. This distribution clearly shows the bias toward frontal regions, with about 70% of the amplitude of the recorded signals in prefrontal and frontal channels being due to EOG artifacts. Further, the proposed model predicted less EOG in the pre-RT period than ICA. The fixation of the participants’ gaze during the period preceding the apparition of a stimulus (see the bottom right panel of Fig. 4 for an illustration of this) suggests that the proposed model distorts the EEG signals less than ICA.

**Fig. 5.**
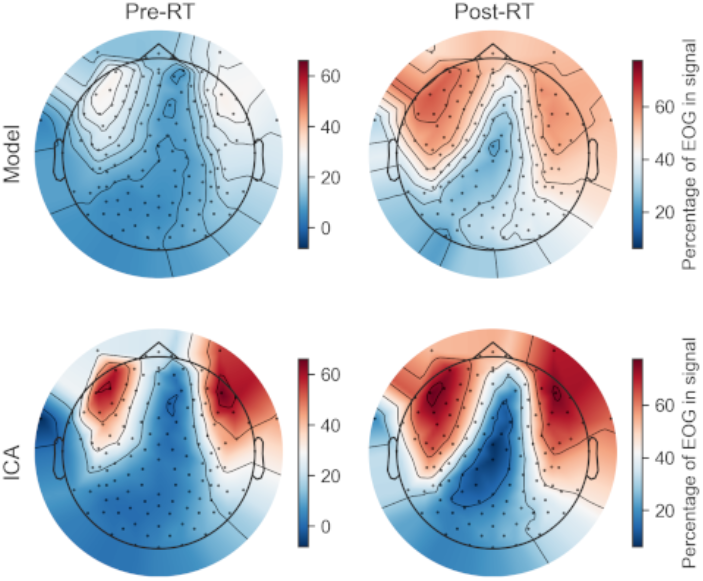
Topographic plots showing the spatial projection of the predicted EOG signal (across participants), as predicted by the model (top row) and by ICA (bottom row). The spatial projection is shown for the pre-RT period (left column) and the post-RT period (right column). We used only trials for dot for this figure.

Next, we used the evoked EEG time series (averaged for each dot) before and after EOG artifact reduction to test the sensitivity and specificity of the proposed deep learning model and ICA. In Fig. 4, we can observe that the deep learning model seems more conservative in EOG reduction than ICA and distorts signals less in the pre-RT period. To quantify the specificity and sensitivity of the proposed model for reducing EOG artifacts, we computed the SNR as described in (4). e averaged SNR values for the pre-RT (specificity) and post-RT (sensitivity) periods within subjects (across runs) and computed a paired t-test to compare the SNR values for the proposed model versus ICA. Fig. 6.a illustrates an example of this approach for channel E25 and dot 25. For this combination of channel and dot, the average values for the Model and ICA indicate a higher specificity for the model but a higher sensitivity for ICA. We repeated this process for all channels and dots and aggregated the result on topomaps for specificity (Fig. 6.b) and sensitivity (Fig. 6.c). The results suggest that this tendency toward higher specificity but lower sensitivity for the model compared to ICA can be generalized across the scalp, except for a higher sensitivity of the model in the posterior region of the scalp.

**Fig. 6.**
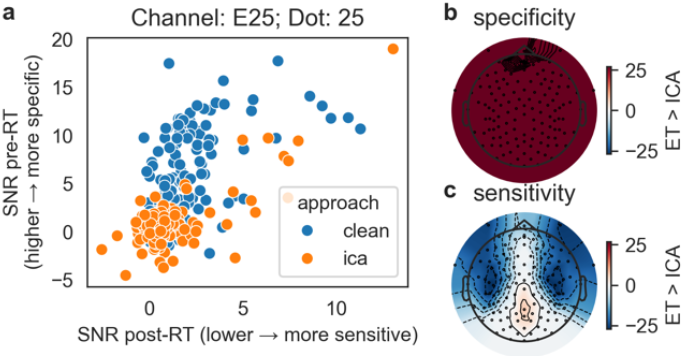
Performance of our approach based on eye-tracking signals (ET) versus ICA. **a)** Example of analysis of sensitivity and specificity for a specific channel (E25, a frontal channel) and dot 25. **b)** Comparative performance of the specificity for the two approaches across the scalp. For each channel and dot, specificity is determined as illustrated in panel **a**. SNR values are then averaged within subjects (across runs), and a paired t-test is computed to compare the SNR values for ET vs ICA. We counted +1 when the specificity was larger for ET than ICA, -1 when it was smaller, and 0 when it was not statistically different (p > 0.05). The sum of these scores is computed across the 27 dot conditions and displayed as a topomap. **c)** Same as for b, but for sensitivity.

## 4. Discussion

This study demonstrated a novel approach for EOG artifact rejection in EEG signals recorded simultaneously with ET. The code implementing this approach is available on GitHub (https://github.com/lina-usc/eog-learn). The emergence and popularization of such recordings [18] has opened new possibilities by providing reference signals closely associated with EOG generation. One of the constant challenges of EOG rejection is the absence of a ground truth for evaluating the effectiveness of proposed approaches. This common situation limits the options available to investigators to 1) using synthesized recordings with a known ground truth but questionable face and ecological validity or 2) using recorded EEG with unknown ground truth. The addition of ET data, although not providing ground truth for EOG artifacts *per se*, partly mitigates this thorny issue by providing reliable information on eye motions usable for inferring EOG artifacts.

Leveraging these signals, we adopted a *data-driven black-box* approach to EOG artifact removal from EEG signals. The *data-driven* qualifier in this statement comes from using ET signals and deep learning to empirically map the association between the movement of the eyes and its impact on EEG signals. We use the *black-box* qualifier to contrast with an approach using ET signals and a physiological model of how eye movements generate EOG artifacts. Our approach does not consider any knowledge specific to the application at hand. The solution relies on the generic task of learning an arbitrary relationship between an input and an output given enough data. This task can be addressed by deep neural networks, which have been shown to work as universal function approximators [19]. Because of the adoption of a generic solution, conceptually, this approach may be suitable for other applications (e.g., removal of electrocardiogram artifacts in EEG, correction for the effect of motion on electrocardiogram signals), as long as the source of the contamination is due to a process for which we have a separate reference signal.

Crucially, the approach we adopted provides an opportunity to assess the performances of blind source separation conducted with ICA more objectively. This technique currently dominates the field [4]. It has been shown to perform well in general, and our analyses support this position in many ways. However, ICA for EOG rejection is known to have limitations [17]. More importantly, although its capacity to remove EOG artifacts can be readily evaluated on noisy EEG signals (e.g., see Fig. 3), the degree to which it may distort neural signals is more difficult to establish. For example, ICA tends to distort the phase of EEG signals [20], [21], a key element in neural dynamics and a property essential for all functional connectivity metrics based on phase consistency (e.g., coherence, phase locking value, phase lag index). Our experiment demonstrated that eye movement information can be used to remove EOG artifacts effectively while distorting neural signals significantly less. Although the lack of a ground truth creates some uncertainty in the interpretation of the SNR-based measures of sensitivity and specificity, our approach has shown a much higher specificity than ICA. In general, ICA has shown a higher sensitivity. However, since we do not have the ground truth for the effect of eye movements on the EEG, we cannot rule out the possibility that the apparent superior sensitivity of ICA in post-RT windows could also be partly due to an over-correction of ICA.

Interestingly, our approach was more sensitive to parts of the scalp that typically are less impacted by EOG artifacts (i.e., central/occipital regions; see Fig. 6.c). Our comparison with ICA relied on the automated classification of independent components associated with eye movement artifacts. The classifier we adopted (i.e., ICLabel) has been designed using machine learning and a large dataset of independent components annotated by experts. This classification is, therefore, vulnerable to biases associated with our understanding of the topological appearance of EOG artifacts. EOG artifacts are known to have the most impact on the frontal region. However, although electrical dipoles generated close to the forehead may have their strongest effect on that part of the scalp, their field wraps around virtually the whole head (with decreasing amplitude due to attenuation). Our results suggest that independent components selected for rejection may tend to undercorrect for these more distant effects. This observation highlights the most significant contribution of this approach: using ET to assess the impact of eye movement on EEG may provide us with a more reliable and objective assessment of EOG artifact topography. We can then use this assessment to develop new methods or correct existing methods that do not require the availability of ET data.

## 5. Limitations

The need for synchronized EEG and ET recordings constitutes the most obvious limitation of the approach we proposed in this paper. However, although we may use this approach directly to clean EEG in such datasets, this application was not the main reason for this study. We would argue that using such a dataset to develop methods that can highlight the limitations of current techniques and suggest possible ways to remedy these shortcomings without requiring ET data is of greater interest. We focused our comparisons on the automated ICA approach because of its popularity for EOG removal.

For our specific application, we attempted to learn the predictable part of the EEG based on ET information. This part is arguably a minor portion of the EEG makeup.

The learning task is, therefore, complicated by the relatively low percentage of predictable information in the signals. We demonstrated that even with a relatively small amount of data (i.e., single recordings), it is possible to learn that relationship with a satisfying degree of precision. However, we could possibly improve the sensitivity and specificity of EOG removal by using a larger training set. The degree to which the performance is saturated with the current size of the training data is unknown.

The data used for this study (i.e., looking at many predefined targets on a screen) were particularly well-suited for our analysis and for learning the mapping between eye movement and EOG artifacts. However, we used information on the structure of the task only for performance analysis. Our training did not rely explicitly on the properties of the experimental protocol (i.e., the whole recording was segmented in 1s epochs and passed to the training routine without any information on the stimuli). However, implicit characteristics (e.g., the systematic coverage of the whole screen area by eye movements) may have been beneficial.

It is also worth considering that the relationship learned between eye movement and the predictable part of the EEG is made up of various contributions, including a component due to the artifact generated by the movement of the eye (the component we generally want to remove) and the neural activity systematically correlated with the eye movement, such as the neural signals controlling the movement of the eye and the neural activity created by the change in visual stimuli when the line of sight shifted. Analysts may or may not want to remove these latter components depending on the hypotheses under investigation. The approach considered in this study cannot disentangle these different components. However, it offers a powerful framework for investigating these components, for example, by using virtual reality to experimentally control changes in the visual field as a function of eye movement.

Lastly, we decided in this study to perform learning and testing on single recordings. One advantage of this approach is that it is constrained to the recordings themselves (e.g., it is independent of the subject sample size). This approach is, therefore, tailored to the specificity of the participants (e.g., the specific shape and electrical properties of the head of the participant affect how electrical currents generated by the movement of the eyes travel and are recorded by the EEG system) and of the recording (i.e., effects of the experimental protocol, environment factors, etc.). Thus, our study did not aim for the generalizability and reusability of these deep learning models. Future work could consider training a single model across a large sample to target generalizability and reusability. It is also possible that by benefiting from a larger sample for training, the mapping learned would be more precise. Whether the gain obtained from training across subjects would offset the loss of precision due to interindividual and within-individual/between-recordings variability is currently unknown. Alternatively, pretrained models may provide an advantageous middle ground.

## 6. Conclusion and Future Works

We presented a deep-learning approach that leverages ET information in synchronized EEG/ET recordings to remove EOG artifacts from EEG recordings. Most importantly, we demonstrated how to harness this objective source of information to benchmark existing approaches and better understand their limitations. In future works, we plan to address the limitations associated with this black-box approach by designing a generative model of how EOG artifacts are generated from eye movements relying on physiological knowledge. Provided that 1) we dispose of enough information to develop a faithful generative model of EOG artifact from measurements of the eyes position and 2) given that precise ET information is available, this source of artifact could be removed accurately from the EEG. Furthermore, in combination with the approach we presented here, the EOG and the neural component associated with eye movements could then be disentangled, allowing more precise analyses of neural activity.

