## Supplementary Materials for "Removing EOG Artifacts from EEG Recordings using Deep Learning"

bioRxiv preprint doi: https://doi.org/10.1101/2024.05.24.595839; this version posted May 26, 2024. The copyright holder for this preprint (which was not certified by peer review) is the author/funder, who has granted bioRxiv a license to display the preprint in perpetuity. It is made available under aCC-BY-NC-ND 4.0 International license.

### Supporting information

#### EEG Sensor Montage

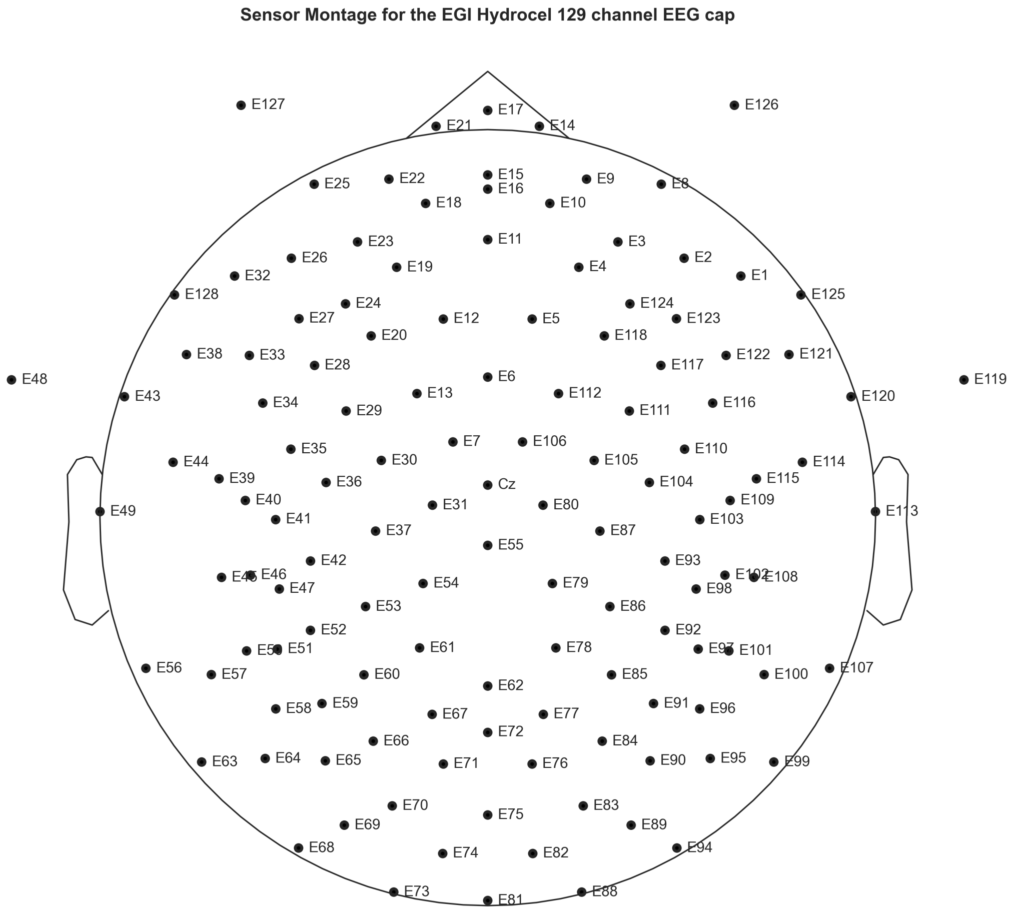


Figure 5: Montage displaying the idealized sensor locations of the EGI 129 channel EEG cap.

bioRxiv preprint doi: https://doi.org/10.1101/2024.05.24.595839; this version posted May 26, 2024. The copyright holder for this preprint (which was not certified by peer review) is the author/funder, who has granted bioRxiv a license to display the preprint in perpetuity. It is made available under aCC-BY-NC-ND 4.0 International license.

#### Eye-tracking Validation

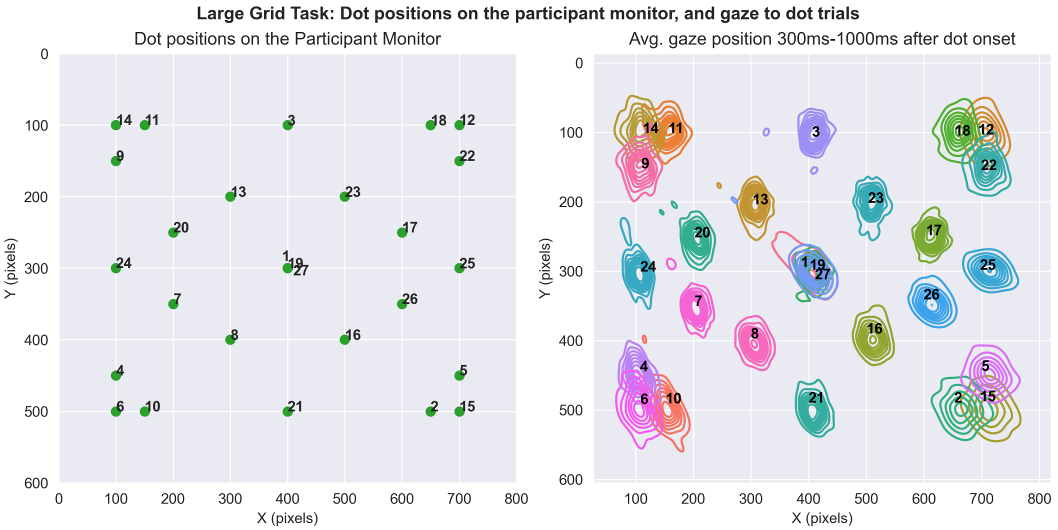


Figure 6: Pixel coordinates on the participant screen where each dot was displayed (left), and the average gaze position in response to the dot onsets, across participants.

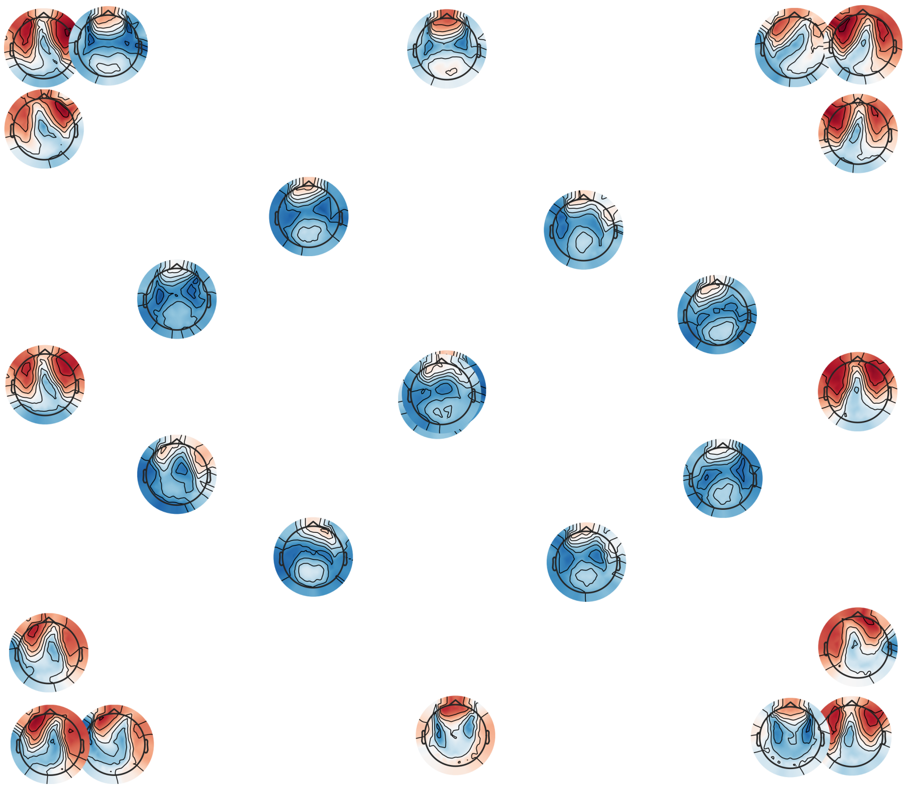

Figure 7: Topographic representation of the predicted EOG after gaze to each dot, averaged across participants.
